# Gene Panel Sequencing in a Chinese High-risk Breast Cancer Cohort

**DOI:** 10.1101/513317

**Authors:** Xianyu Zhang, Xiaohong Wang, Bingbing Song, Kang Shao, Guibo Li, Wenjing Jian, Cong Lin, Min Wang, Xiaofei Ye, Jingjing Xie, Bingshu Xia, Shouping Xu, Boyang Cao, Liyun Xiao, Zhao Zhang, Meng Liu, Enhao Fang, Haoxuan Jin, Xiaofeng Wei, Michael Dean, Jian Wang, Huanming Yang, Xianming Wang, Shida Zhu, Yong Hou, Da Pang

## Abstract

Currently, over 20 genes have been defined that can confer susceptibility for high-risk breast cancer. Although research has proved the utility of multiple-gene sequencing in the assessment of breast cancer risk, there is little data from China patients. Here, we use a multiple-gene sequencing panel to identify the variant spectrum in Chinese high-risk breast cancer subjects.

A total of 829 Chinese high-risk breast cancer patients participated in the research. The coding regions of 115 hereditary cancer susceptibility genes were sequenced using a next generation sequencing platform. In total, 193 pathogenic variants were identified in 45 genes from 177 patients. The pathogenic variant carrier rate is 21.4%: with 10.5% patients carrying a BRCA1 or BRCA2 mutation only, 10.0% of patients carried non-BRCA gene mutations only, while 1.0% of patients carried both a BRCA1/2 and a non-BRCA gene mutation. Variants of uncertain significance (VUS) totaling 2632 were identified in 115 genes from 787 of 829 patients: 82.5% patients carried more than one VUS, and only 5.1% patients did not carry any VUS. Families carrying pathogenic variants were tracked and adenoma was founded in three of them. Our data provide a comprehensive analysis of potential susceptibility variations of high-risk for breast cancer in a Chinese population. This data will be useful for the comparison of the susceptibility variation spectrum between different populations and to discover potential pathogenic variants to improve the prevention and treatment of high-risk breast cancer.

## Introduction

Breast cancer is the most common cancer in women worldwide with over 1.6 million new cases diagnosed and over 520,000 deaths annually (Globocan 2012, http://globocan.iarc.fr/Pages/fact_sheets_cancer.aspx). Improved treatment has contributed to steady declines in breast cancer mortality in developed countries (1). However, breast cancer incidence is continuously rising in nearly all countries and mortality rate stays high in most Asian countries (Globocan 2012). In China, there is an estimated 272,000 total cases (269,000 in women) and 70,000 total deaths annually (69,000 in women) (2).

Up to 10% of breast cancer is caused by the inheritance of germline mutations in susceptibility genes (3). Among them, BRCA1 and BRCA2 are the predominate genes, while other genes such as PALB2, TP53, PTEN may also contribute to genetic risk, and screening susceptible population for such mutations can lead to reduced mortalities in breast and ovarian cancer (4–7). There is a considerable diversity in mutations in BRCA1, BRCA2 and other genes in different populations, requiring studies worldwide (3, 8–11). Studies of germline mutation spectrum have been carried out in Asian populations, but comparatively few in Chinese populations (12–18). This study aims to identify the spectrum of germline variants in a large panel of cancer genes in selected Chinese breast cancer families.

## Method

### Participants and Selection Criteria

A total of 829 breast cancer patients who met genetic risk evaluation criteria according to National Comprehensive Cancer Network (NCCN) guidelines for breast and/or ovarian cancer genetic assessment (version1.2014) were recruited from 2010 to 2016. All patients met the high risk criteria: 1. early age-of-onset, people suffer breast cancer under age 45; 2.patients who have at least two primary breast cancer; 3.with a family history: whose first degree relatives have breast cancer or ovarian cancer. Of the 829 breast cancer patients, 593 came from Harbin Medical University Cancer Hospital, and 236 came from Shenzhen Second People’s Hospital of China. All of them did not have previous breast cancer susceptibility gene testing.

### Ethics, consent and permissions

Every participant signed an institutional review board-approved informed consent document offered by Harbin Medical University Cancer Hospital, Shenzhen Second People’s Hospital or BGI Shenzhen. The consent informed the participants that their test data would be used for research. Clinical data and family disease history information were including patient gender, age at diagnosis of breast cancer, breast cancer molecular subtype, site, personal history of other cancers, and family history of cancer in close blood relatives: diagnosis age, tumor type, and health status. The clinical information is shown in Supplementary Table S1.

### Gene Panel

All breast cancer samples were subjected to the target sequencing using a multiple-gene panel and subsequent variant analysis. A total of 115 target genes were selected through a review of databases (HGMD: Human Gene Mutation Database, NCBI ClinVar database) or published articles on the role of the genes in hereditary cancer. The main types of hereditary cancer include Breast cancer, Colorectal cancer, Gastric cancer, Prostate cancer, Thyroid cancer, Renal cancer and other cancers which have a genetic risk. The Breast cancer susceptibility genes (27 genes) are shown in Table S2; the other 88 cancer susceptibility genes involved in 35 cancer types are shown in Supplementary Table S2. The panel has been used for previous research (19), but is not currently commercially available. Interested individuals can contact us for more details.

### Sample Treatment, Next-Generation Sequencing and Variants Calling

Sample preparation and DNA sequencing were performed at BGI Shenzhen. The samples were separated into two groups and analyzed on different sequencing platforms. A total of 634 samples from Harbin Medical University Cancer Hospital or Shenzhen were sequenced on the Blackbird platform (Complete Genomics, a BGI Company). An additional 195 samples from Shenzhen were sequenced on the Hiseq 2500 platform (Illumina) with the Paired-end 91 bp strategy.

DNA was extracted from participant’s peripheral blood sample by Qiagen DNA Blood mini kit (Qiagen, Hilden, Germany) according to the manufacturer instructions. Qubit Fluorometer (Life Technologies) and agarose gel electrophoresis were used to detect DNA concentration and integrity.

For the samples on Blackbird platform, the DNA amount used for library construction was 1ug. DNA was randomly fragmented to 200-400bp and the A-adaptor was added. The coding region and coding region ±30 boundaries of 115 genes were captured by a BGI capture array (produced by BGI). Double strand DNA was cyclized after the PCR amplification. Then the B-adaptor was added, and the sample converted to a circular single-stranded DNA. After rolling circle replication, the DNB (DNA nanoball) was loaded onto the chip and subsequent DNA sequencing was performed according to published protocols to at least an average depth of 390X and 99% coverage on target regions [38]. Over 0.6G base of data was generated for every sample. Variants were detected using Small Variant Assembler Methods which was available on the CG website (http://www.completegenomics.com/documents/Small_Variant_Assembler_Methods.pdf). Next, variants were filtered by allele depth, allele frequency, mapping quality and region of variation (major parameters set as follows: alteration allele > 1, Allele depth > 8, BAF > 30%, not in highly repeat region, no Indels in 10bp and other parameters set to default).

For the samples on the Hiseq platform, genomic DNA with initial amount above 1ug was randomly fragmented to 200-300bp by Covaris E210 (Massachusetts, USA). Then the library was constructed as follows: end-repair, A-tailing, adapter ligation, and PCR amplification. PCR products were captured by the same BGI chip as in Blackbird platform. Then quantified by quantitative PCR and pooled for sequencing on the Hiseq 2500 (Illumina) according to the manufacturer’s protocols (https://support.illumina.com/content/dam/illumina-support/documents/documentation/system_documentation/hiseq2500. Over 0.6G base data was generated for every sample with an average depth of about 200X and over 99% coverage on target regions. Reads were filtered by SOAPnuke1.5.0 with parameters -l 5 -q 0.5 -n 0.1and assembled by BWA 0.7.12. Samtools 1.2 and picard MarkDuplicates 1.138 with standard parameters used in Bam file processing and duplication marking. The base quality recalibration and local realignment were performed by GATK 3.4. Variants were called by GATK 3.4 and further filtered by quality depth, strand bias, mapping quality and reads position (major parameters were -filter “QD < 2.0 ‖ FS > 60.0 ‖ MQ < 40.0 ‖ SOR > 4.0 ‖ MQRankSum < −12.5 ‖ ReadPosRankSum < −8.0” for SNP_filter, and −filter “QD < 2.0 ‖ FS > 200.0 ‖ ReadPosRankSum < −20.0 ‖ SOR > 10.0” for INDEL_filter).

### Variant Classification

Variants were annotated by ANNOVAR and then classified as pathogenic, variants of uncertain significance (VUS) and benign according to American College of Medical Genetics (ACMG) recommendations(20, 21) by a semi-automatic pipeline called VCE for interpretation of sequence variants. The detailed evidence can be found in Supplementary Table S4. Most pathogenic variants detected by next-generation sequencing were confirmed by conventional PCR-Sanger sequencing, except variants located in repeat regions.

### Gastroscopy of mutation carrier

Mutation carrier families accepted the clinical screening such as gastroscopy, fibercolonoscopy, breast ultrasound and serum tumor marker detection (CEA, CA199, CA742) based on the hereditary cancer susceptibility gene research progress. Up to now, gastroenteric precancerous lesions have been observed in 3 families with ATM, PMS2, and PALB2 pathogenic variants respectively.

### Functional analysis of Variant of uncertain significance

Functional analysis of variants from BRCA genes was conducted by Ranomics Inc. To validate variants occurred in RING domain, they measured if mutated BRCA1 showed decreased ligase activity and weaker protein-protein interactions in its heterodimer with the BARD1 protein(22) They also validated variants in BRCT domain by comparing growth speed between dysfunctional variants and normally functioning variants in the BRCA1 BRCT domain(23).

### Statistical analysis

The statistics in this research is performed by R 3.5.1. The significance between mutation prevalence and characteristics is compared by Chi square test or Fisher exact test depending on case number.

## Results

### Variants classification pipeline

The panel we used in this research contains a high number of genes, leading to a low efficiency in classification by manual analysis. We consequently developed a semi-automatic pipeline called Variant Clinical Explanation (VCE) for clinical classification of germline variants related with cancer by ACMG/AMP 2015 guideline. VCE combine automatic classification and manual check referring to the method of InterVar(24), which would judge each variant with all the evidence and assign it to a rough class. Then apply manual check evidence to some variants. VCE changed the gene set, supporting databases and filter criteria. Compared with another variants pathogenicity interpreting and predicting software CharGer (Characterization of Germline variants)(25), VCE’s difference lies in more gene number, more support database and silicon prediction software and hints for variants that need manual review. Therefore, it is suitable for border gene panel analysis and could provide more assistant information for manual review (Table 1). In total, the variants need to be manual checked is about 10%.

**Table 1.**
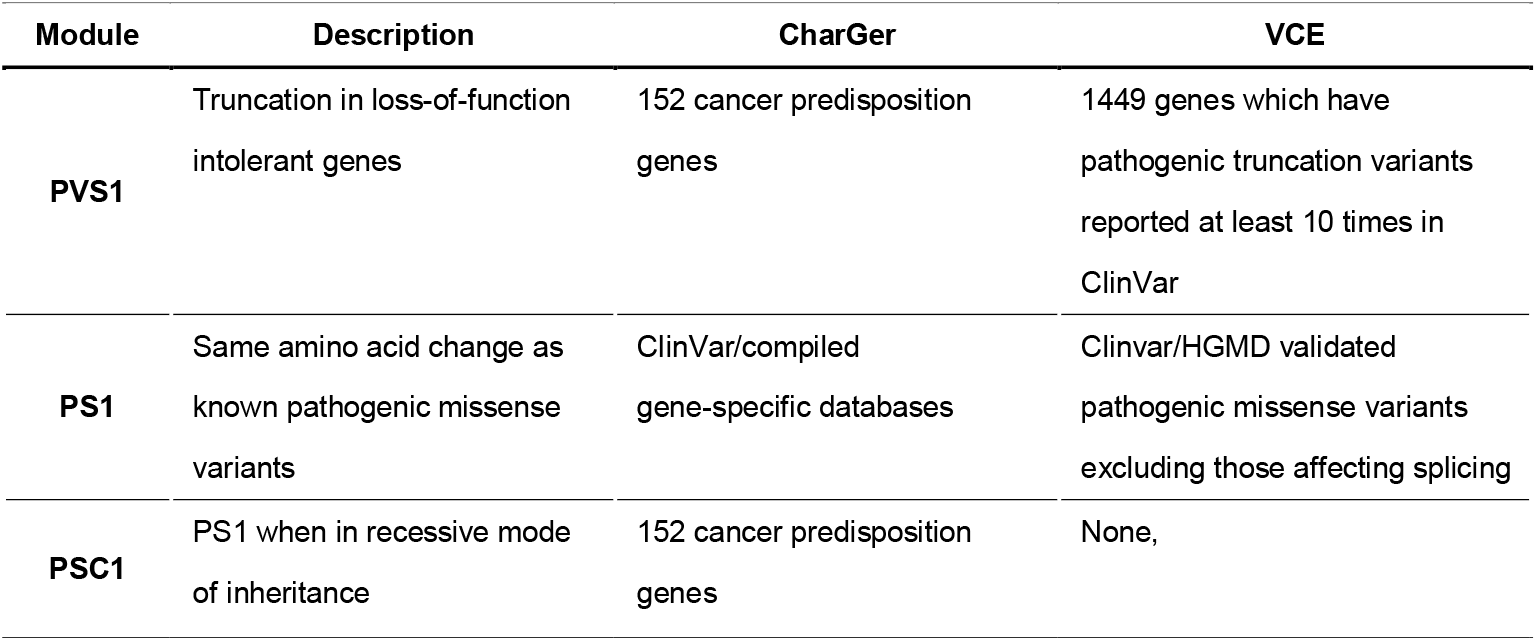
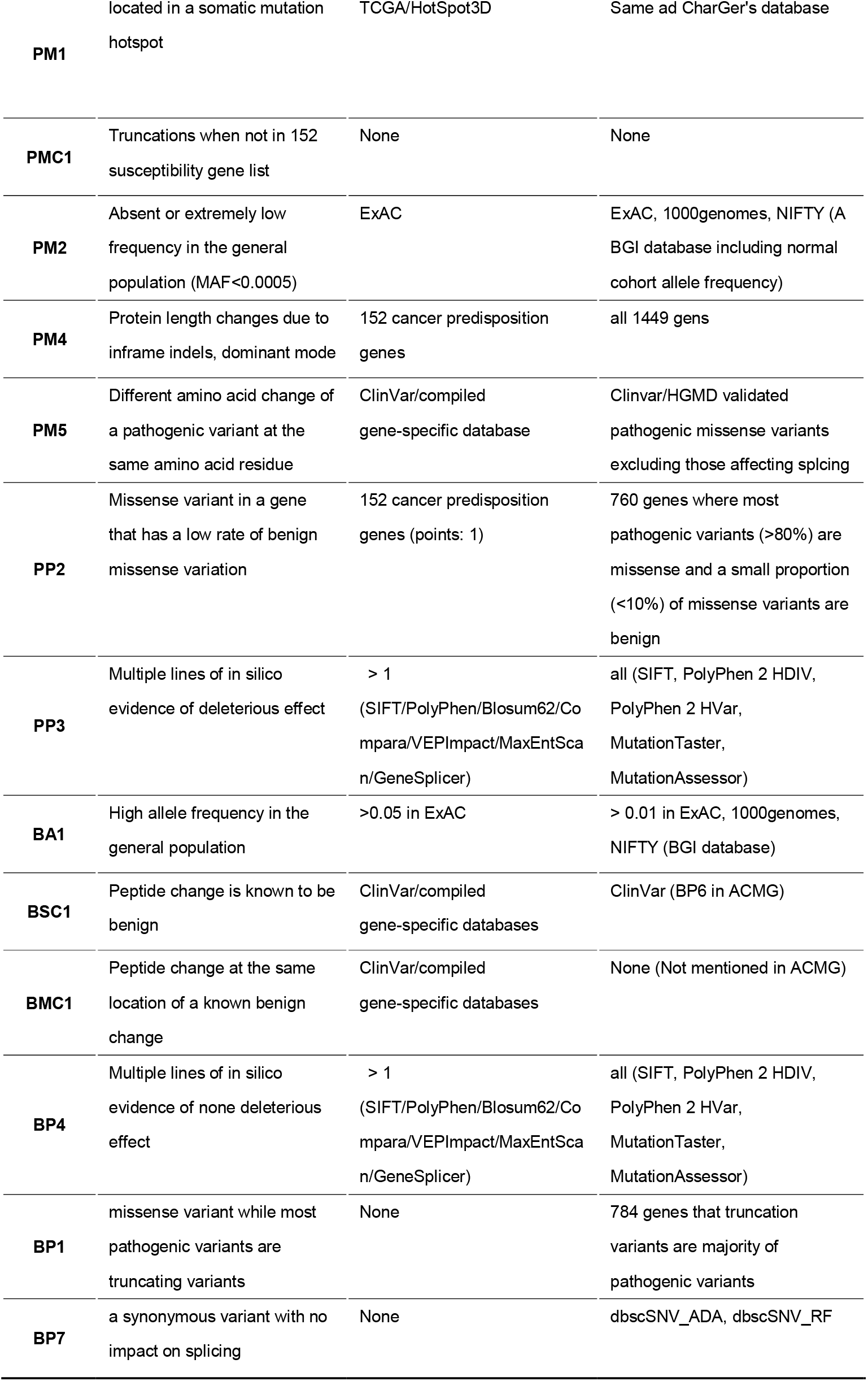
The table below is the different interpretations on evidence between VCE and CharGer. VCE could apply to more genes and different database.

### Total Distribution of Pathogenic Variants

Mutation analysis of 115 susceptibility gene for hereditary cancer was performed in 829 breast cancer patients from Northeast part of China or Shenzhen city of China. In total, 193 pathogenic variants were identified in 45 genes from 174 different patients (Fig1 A), and the pathogenic variant carrier rate is 21.4%: with 87 (10.4%) patients carrying a *BRCA1 or BRCA2* mutation, 38 (4.6%) patients carried other breast cancer susceptibility gene mutations, 38 (4.6%) patients carried other cancer susceptibility gene mutations, and 12 (1.5%) patients carried both a *BRCA1/2* and other BC susceptibility gene mutation (Fig1 B). Among those variants in breast cancer susceptibility genes, 11 recurrent (detected in more than one patient) pathogenic variants were identified in 5 genes: 4 in *BRCA1*, 5 in *BRCA2*, 1 in *HMMR, PALB2* and *PTEN* respectively. Eight recurrent pathogenic variants were identified in 7 genes: 2 in *ZFHX3*, 1 in *FANCG, MPL, PRF1, NF1*, *NTRK1*, and *SBDS* respectively. Detailed information can be found in Supplementary Table S4.

**Figure 1A.**
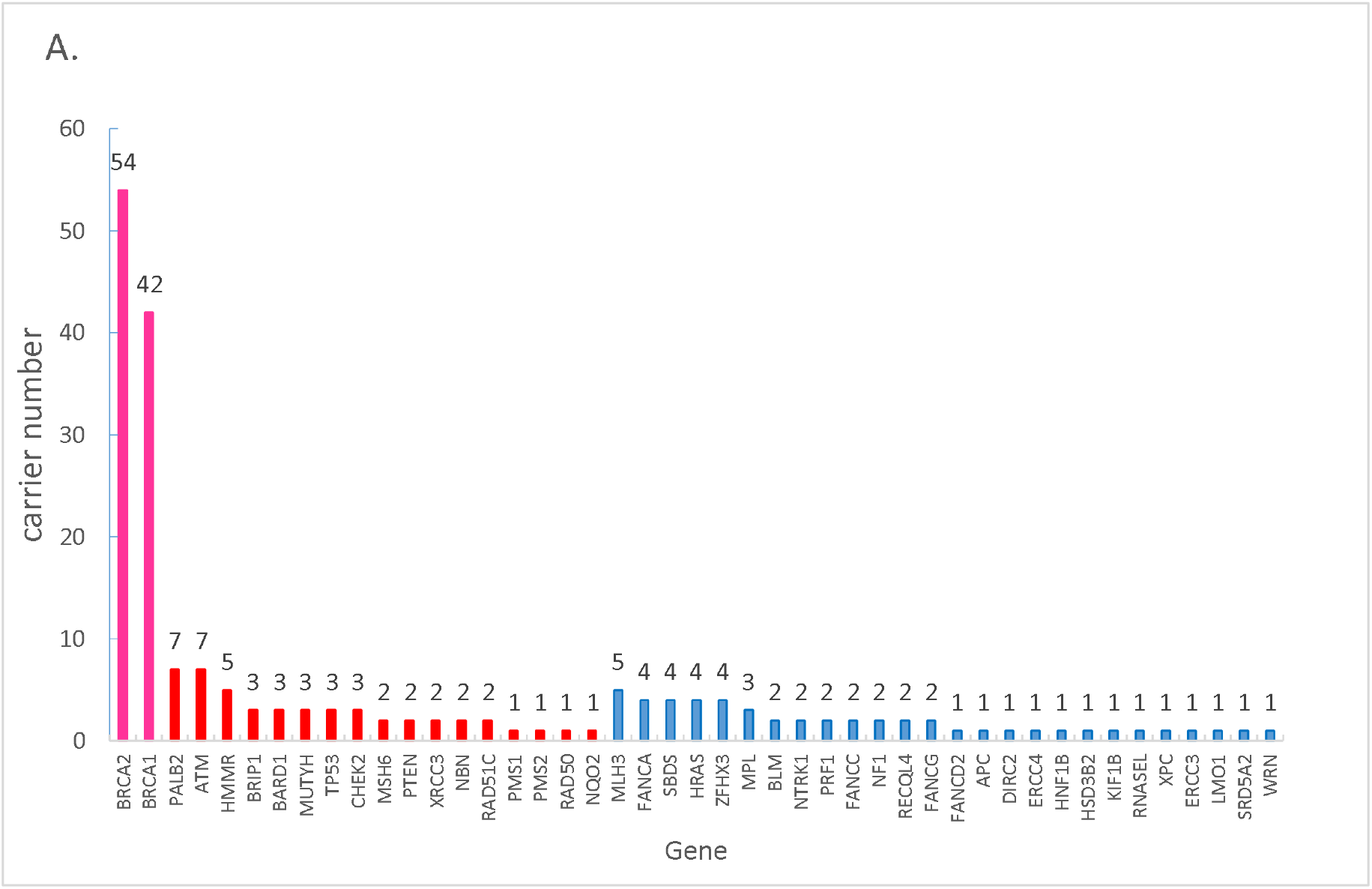
The pathogenic distribution of BRCA1/2 (pink), breast cancer risk related other BC susceptibility genes (red) and other cancer susceptibility genes (blue).

**Figure 1B.**
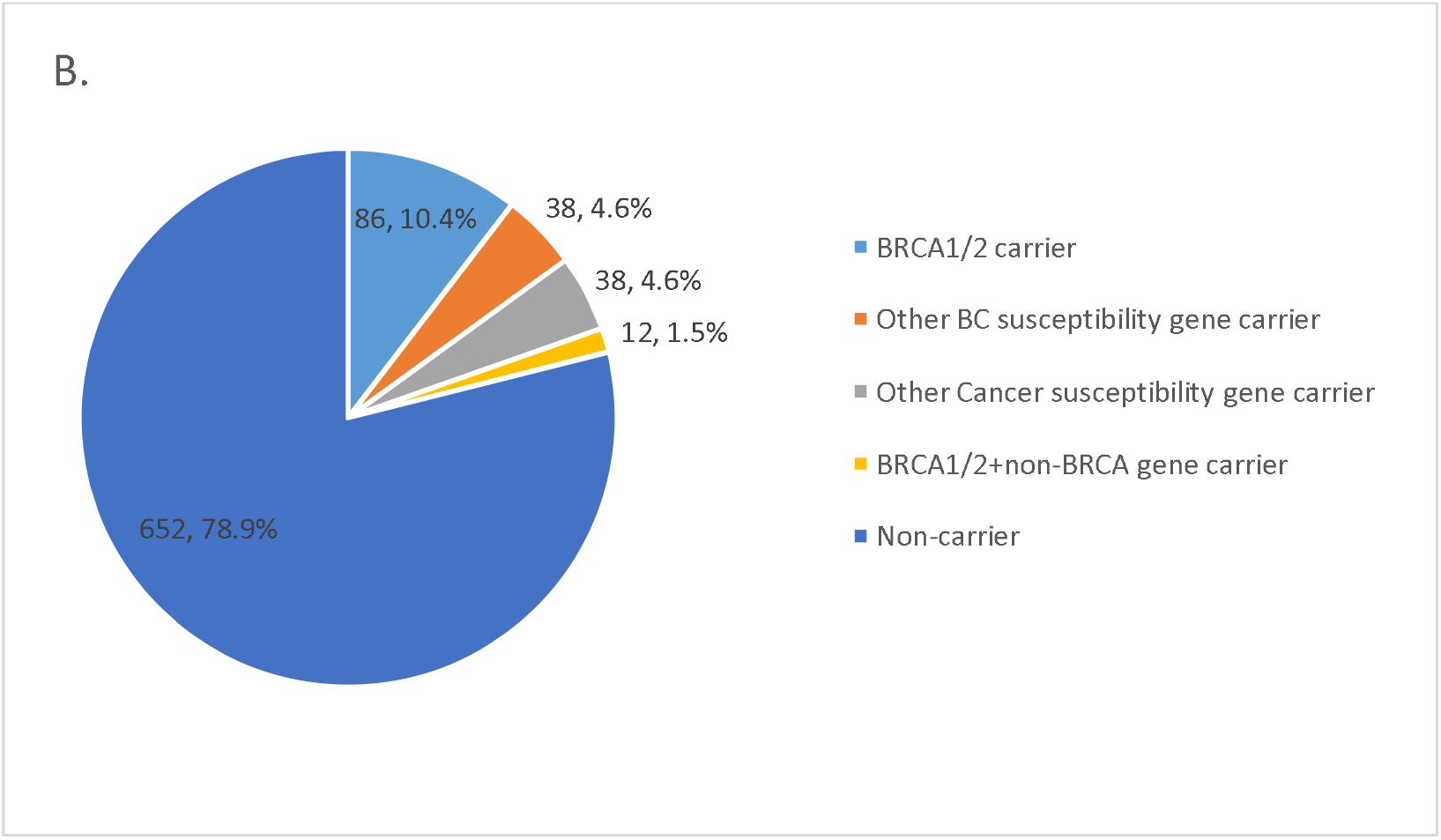
The pathogenic variants distribution in the cohort. Variants were classified by 4 main groups. BRCA1/2 group, other breast cancer susceptibility gene group, other cancer susceptibility gene group, and non carrier. There is also a small proportion of people carried more than one variants that belongs to different groups.

### *BRCA1* and *BRCA2* Pathogenic Variants and Clinical characteristic

The prevalence of *BRCA1* and *BRCA2* pathogenic variant in 829 breast cancer patients is 5.0% and 6.5% respectively. The *BRCA1/2* mutation frequency found in our research is similar with the results from other studies with Chinese population, but distinct from reports based on other populations (Table2).

**Table 2.**
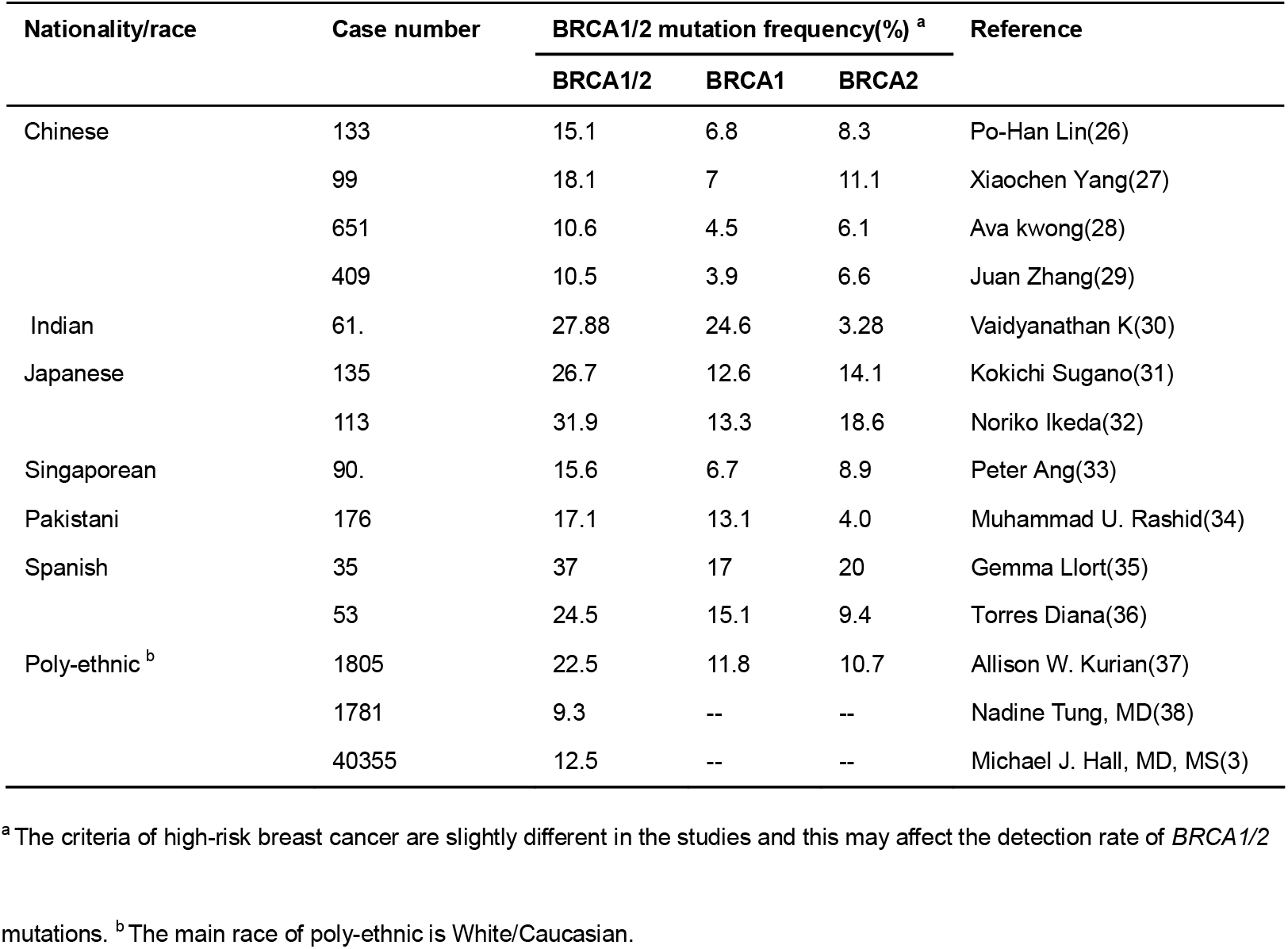
*BRCA1* and *BRCA2* mutation frequency in high-risk breast cancer patients of different nationality/race.

**Table 3.** Characteristics of patient information and comparison of BRCA1/2 mutation prevalence between different characteristic. P1 means p-value of BRCA1 carriers VS. non-carriers, P2 means p-value of BRCA2 carriers VS. non-carriers, P3 means p-value of BRCA1 carriers VS. BRCA2 carriers. *: families with BC compared with families with no Cancer

|  | Basic info | Case | BRCA1 | BRCA2 | PALB2 | ATM | P1 | P2 | P3 |
| --- | --- | --- | --- | --- | --- | --- | --- | --- | --- |
|  |  | num(ratio) |  |  |  |  |  |  |  |
| Gender | Male | 14 (1.7%) | 1 | 4 | 0 | 1 | 0.5 | 0.01 | 0.39 |
|  | Female | 815 (98.3%) | 39 | 50 | 7 | 6 |  |  |  |
| Diagnostic age | 20-29 | 61 (7.3%) | 4 | 3 | 1 | 0 | 0.31 | 0.093 | 0.73 |
|  | 30-39 | 325 (39.2%) | 12 | 16 | 1 | 2 |  |  |  |
|  | 40-49 | 276 (33.3%) | 16 | 18 | 3 | 4 |  |  |  |
|  | ≥ 50 | 146 (17.6%) | 8 | 15 | 2 | 1 |  |  |  |
|  | Unknown | 21 (2.6%) | 0 | 2 | 0 | 0 |  |  |  |
| Breast cancer Pos | Unilateral | 747 (90.1%) | 33 | 46 | 7 | 6 | 0.12 | 0.3 | 0.76 |
|  | Bilateral | 66 (8.0 %) | 6 | 6 | 0 | 0 |  |  |  |
|  | Unknown | 16 (1.9%) | 1 | 2 | 0 | 1 |  |  |  |
| Molecular subtype | Luminal A | 209 (25.2%) | 6 | 11 | 2 | 3 |  |  |  |
|  | Luminal B | 245 (29.6%) | 7 | 20 | 2 | 1 |  |  |  |
|  | HER2+ | 66 (8.0%) | 2 | 1 | 0 | 0 |  |  |  |
|  | TNBC | 146 (17.6%) | 19 | 9 | 1 | 1 | <0.001 | 0.96 | 0.002 |
|  | Unknow | 163 (19.6%) | 6 | 13 | 2 | 2 |  |  |  |
| Family | with BC* | 267 (32.2%) | 17 | 32 | 5 | 4 | 0.027 | <0.001 | 0.562 |
| member | with other C | 174 (21.0%) | 13 | 8 | 0 | 1 | 0.013 | 0.64 | 0.175 |
|  | with no C | 376 (45.4%) | 10 | 14 | 2 | 2 |  |  |  |
|  | Unknown | 12 (1.4%) | 0 | 0 | 0 | 0 |  |  |  |
| Total |  | 829 (100%) | 40 | 54 | 7 | 7 |  |  |  |

**Table 4.** Pathogenic variation of other cancer genes

| Gene | Carrier Num. | Cancer type |
| --- | --- | --- |
| **MLH3** | 5 | Colorectal Cancer(47) |
| **FANCA\*** | 4 | AML, ALL(48-50) |
| **SBDS\*** | 4 | MDS, AML(51, 52) |
| **MPL\*** | 3 | MDS, AML(53, 54) |
| **BLM\*** | 2 | ALL(55, 56) |
| **NTRK1\*** | 2 | Thyroid Cancer(57) |
| **PRF1** | 2 | Lymphoma, Leukemia(58-60) |
| **FANCC\*** | 2 | AML, ALL(49) |
| **NF1** | 2 | neurofibroma, glioma(61, 62) |
| **RECQL4\*** | 2 | Osteosarcoma, AML(63, 64) |
| **FANCG\*** | 2 | AML, leukemia(49) |
| **HRAS** | 2 | neuroblastoma, bladder cancer(65) |
| **ZFHX3** | 2 | Prostate(66) |

\* Means genetic defects in this gene will result in autosomal recessive disorder.
**AML, acute myeloid leukemia; ALL, acute lymphoblastic leukemia; MDS, myelodysplastic syndrome**

We found 9 recurrent pathogenic variants in *BRCA1* and *BRCA2:* 4 in *BRCA1* (NM_007294:c.1465G>T, NM_007294:c.3294delT, NM_007294:c.5156delT and NM_007294:c.5470_5477delATTGGGCA) and 5 in *BRCA2* (NM_000059:c.5864C>G, NM_000059:c.6698_6699insTTTT, NM_000059:c.7617+1G>A, NM_000059:c.9070_9073delAACA and NM_000059:c.9382C>T). *BRCA1* NM_007294:c.1465G>T and *BRCA2* NM_000059:c.5864C>G were previously reported in Chinese high-risk breast cancer(39–42), and another variant *BRCA1* NM_007294:c.5470_5477delATTGGGCA was reported several times in Asian populations(42, 43). However, these variants have not been reported in other ethnic groups. *BRCA2* NM_000059:c.7617+1G>A and BRCA2 NM_000059: c.9382C>T were previously reported in European subjects(44–46). The remaining four (*BRCA1* NM_007294:c.3294delT, *BRCA1* NM_007294:c.5156delT, *BRCA2* NM_000059:c.6698_6699insTTTT and *BRCA2* NM_000059:c.9070_9073delAACA) are novel variants, these variants need to be further validated in large-scale studies.

Among enrolled 829 patients,815 (98.3%) of them are female. 386 (46.5%) of the females are younger than age 40, 422 (50.9%) are older than age 40. The age information of the rest 21 patients (2.6%) is missing. The number of BRCA1/2 carrier is not significantly high in the younger group, but BRCA1/2 carrier number is significantly higher in patient with BC family history than the patient without BC family history and BRCA1 carrier is significantly high in triple negative breast cancer patients than non-triple negative breast cancer patients.

### Non-BRCA Gene Pathogenic Variants

In all, 98 pathogenic variants were identified in 43 non-BRCA genes from 90 patients, the prevalence of non-BRCA gene variants in 829 patients is 10.9%. The number of variant loci is 82, and 18 variant loci were reported in the NCBI ClinVar database or published literature, the rest of the 64 (78%) variants is novel. Detailed information can be found in Supplementary Table S4. In addition, 11 variant loci appeared two or more times, including *FANCG* NM_004629:c.572T>G, *HMMR* NM_012484:c.1989_1990insA, *MPL* NM_005373:c.1908A>G, *NF1* NM_001128147:c.1736_1737insT, *NTRK1* NM_001007792:c.627+2T>C, *PALB2* NM_024675:c.2167_2168delAT, *PRF1* NM_005041:c.65delC, *PTEN* NM_000314:c.697C>T, *SBDS* NM_016038:c.258+2T>C, *ZFHX3* NM_001164766:c.6842_6843insT and *ZFHX3* NM_001164766:c.6847_6848insAG. There were 44 variants in the 18 genes (40.9%, including *ATM, BARD1, BRIP1, CHEK2, HMMR, MLH3, MSH6, MUTY, NBN, NQO2, PALB2, PMS1, PMS2, PTEN, RAD50, RAD51C, TP53* and *XRCC3*) that have been reported in high-risk breast cancer studies, the remaining 26 (59.1%) genes need to be further investigated and validated in high-risk breast cancer diseases.

### Clinical screening of mutation carrier

Families which carry pathogenic mutations in the genes other than BRCA1, BRCA2 an PALB2 received further clinical screening as described in method section. Pre-cancer lesions were found in three families. In the first family, one PMS2 pathogenic variant carrier (proband, study ID: 30033, diagnosed with breast cancer at 55 years old) was found to have colon polyp (diagnosed at 58 years old). Her sister (study ID: 30033FM1), who is also a breast cancer patient (diagnosed at 44 years old), was found to have colon villioustublar adenoma with low-grad intraepithelial neoplasia (diagnosed at 52 years old) and the lesion was removed by endoscopic resection (Fig 2A). They were also both diagnosed with stomach fundus polyp and gastritis. Their father and father’s sister already died because of stomach cancer. They were recommended to do examination again one year later. In the second family, one ATM gene mutation carrier was diagnosed with breast cancer at age 81, (proband study ID30491) who refused the clinical screening because of advanced age. He has two daughters who also carry ATM gene mutation. One of them (study ID: 30491FM2) had been diagnosed with breast cancer (diagnosed at 39 years old), the other one (study ID: 30491FM3) was found to have colon tubular adenoma, atrophic gastritis and stomach fundus erosion in this screening (at 58 years old) (Fig 2B). In the third family, the proband (study ID: 30306) carries both ATM and PALB2 mutation (diagnosed with breast cancer at 47 years old). She was found to have colon villioustublar adenoma and superficial gastritis with erosion (diagnosed at 48 years old) (Fig 2C). Her niece (study ID: 30306FM2) was found to have rectal tubular adenoma (diagnosed at 39 years old), however, she is not a mutation carrier. All adenoma was removed during colonoscopy.

**Figure 2.**
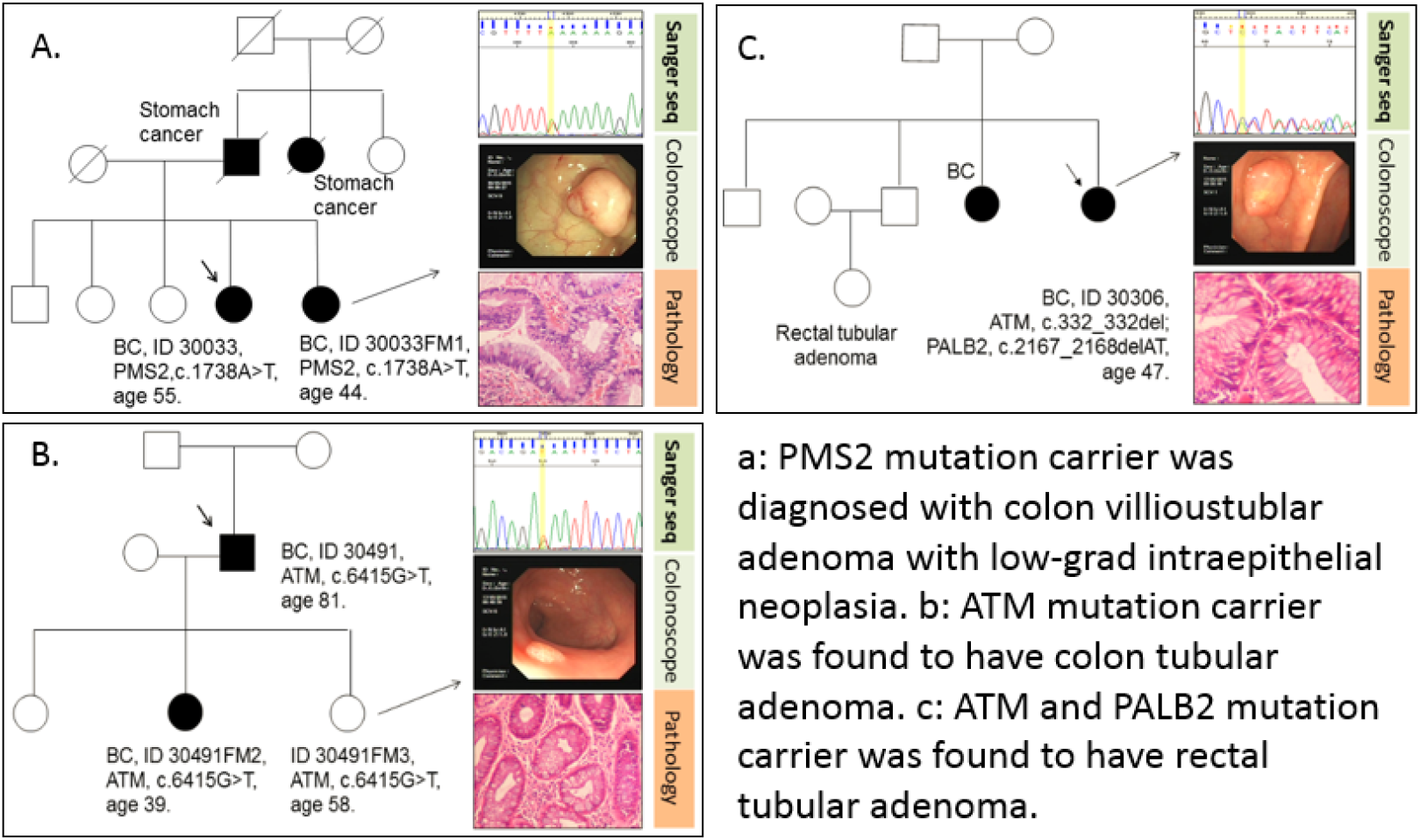
The Clinical screening of three pathogenic variants carrier families

### VUS distribution

All VUS in 115 hereditary cancer susceptibility genes were detected and analyzed. In total, 2632 VUS were identified in 115 genes from 787 of 829 patients. The total VUS number in 27 breast cancer susceptibility genes was 708 (Fig3 A), of which *BRCA1* and *BRCA2* had 5.1% and 12.1% VUS respectively. Meanwhile, we analyzed the VUS frequency distribution in all patients and found that 684 (82.5%) patients carried more than one VUS, and 42 patients did not carry any VUS (Fig3 B). Detailed information can be found in Supplementary Table S5.

**Figure 3A.**
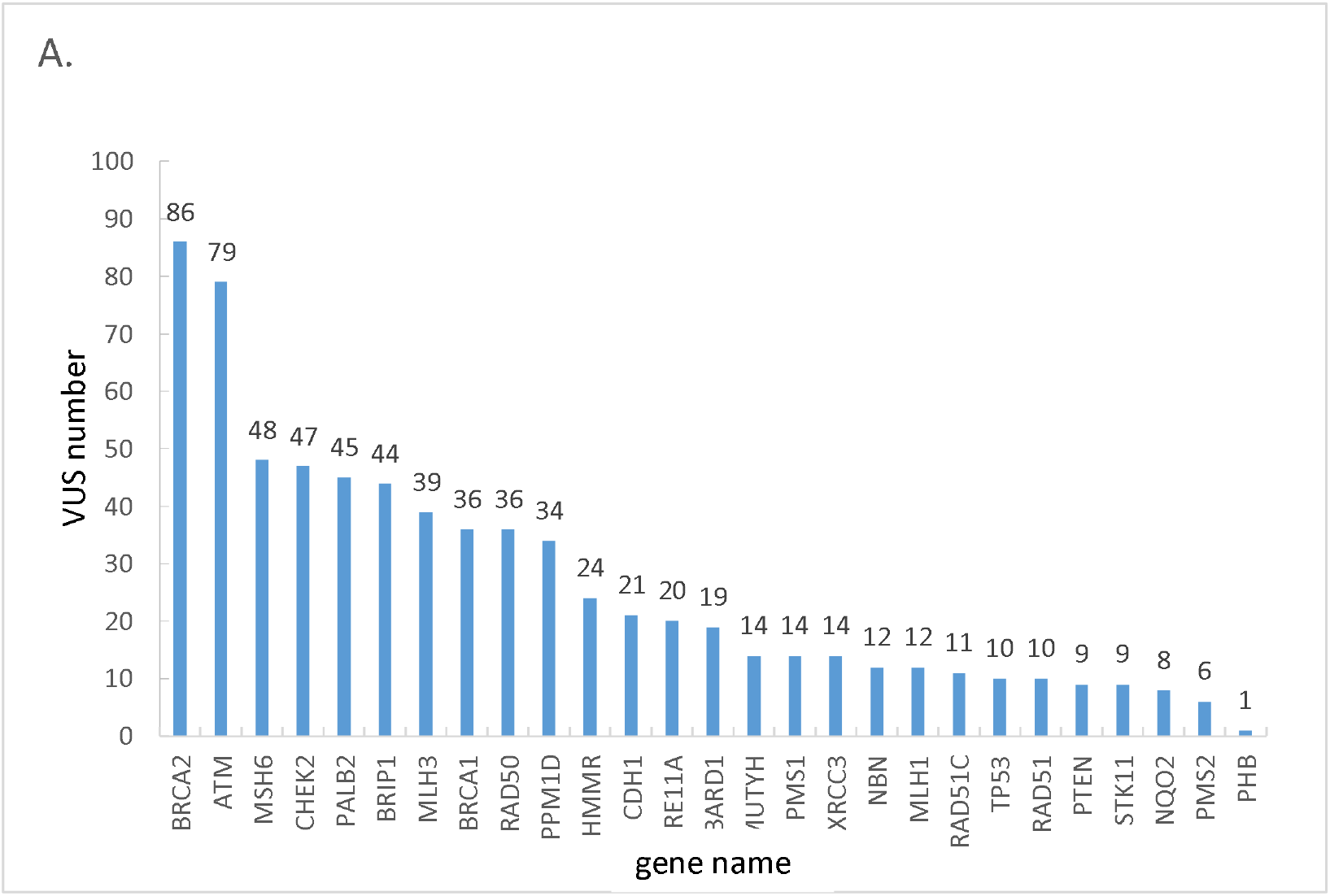
Total VUS distribution in genes.

**Figure 3B.**
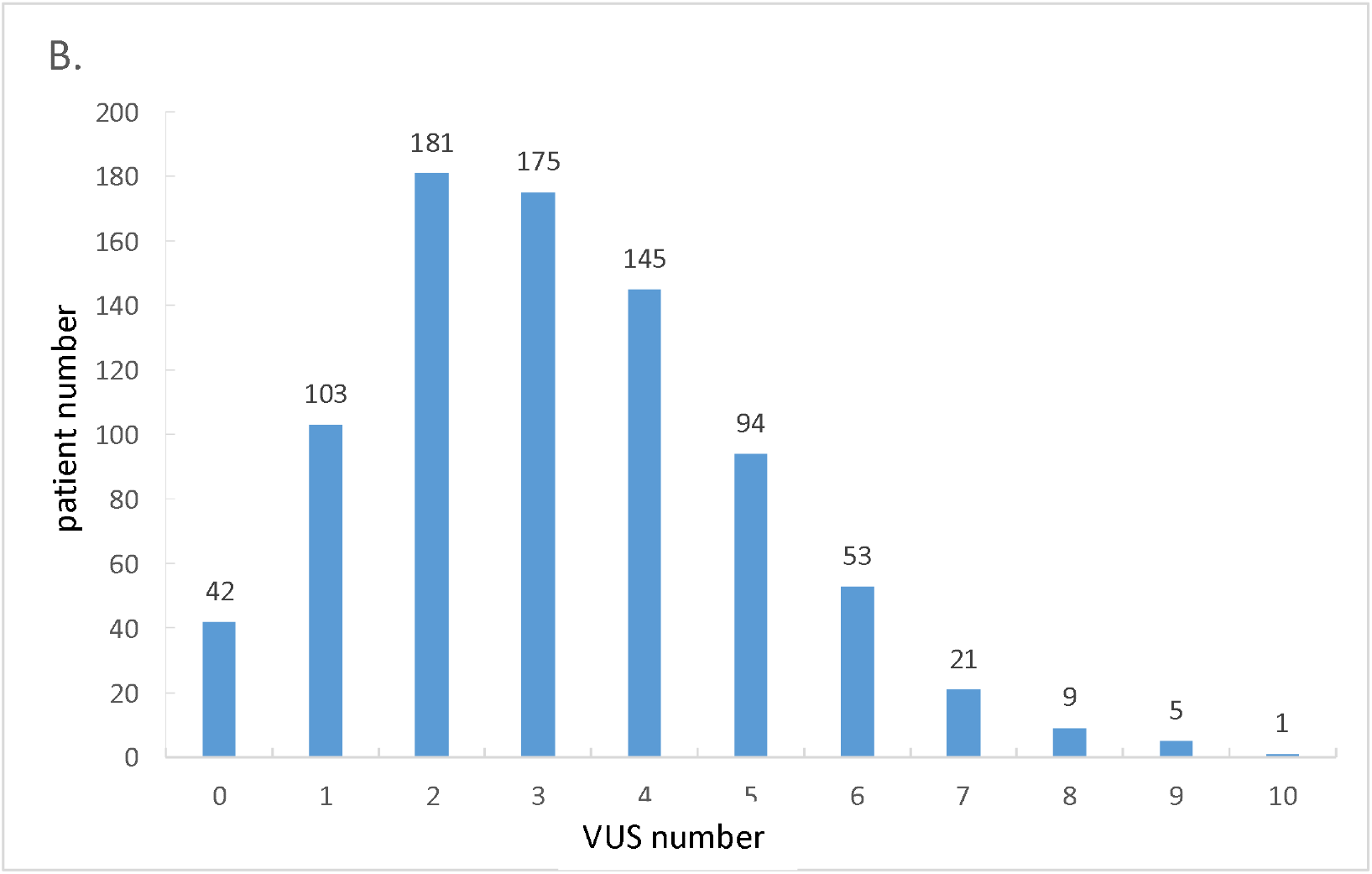
The average VUS number distribution in VUS carriers.

### Functional validation of VUS

All 5 VUS in BRCA1 RING domain and BRCT domain were (seeing in table 5) submitted to a company for functional validation, which was conducted with biochemical function assay (RING domain) and phenotype recapitulation assay (BRCT domain).

**Table 5.** VUS functional validation results

| Gene variant | RING/BRCT domain in BRCA1 | Classification |
| --- | --- | --- |
| p.Cys24Arg | RING | Pathogenic |
| p.Leu1679Val | BRCT | Benign |
| p.Val1809Phe | BRCT | Benign |
| p.Ile1807Val | BRCT | Benign |
| p.Leu1679Val | BRCT | Benign |

One variants (p.Cys24Arg) was defined as pathogenic, and Other 4 variants were defined as benign. The result is consistent with recently reported massive BRCA1 functional analysis(67).

## Discussion

The major hurdle in utilizing NGS data lies in how to interpret the genotype-phenotype relationships, especially in clinical settings. The first key element for resolving the issue is automatic analysis: A large number of germline variants are called after WES/WGS sequencing, it will be time-consuming if we completely depend on manual judgement. Secondly, the standardization: A high proportion of variant classifications is discrepant between intra- and inter-laboratory settings which can be attributed to a different understanding on ACMG guideline and lack of unified standard(68). Thus, the experts review based on comprehensive evidence is also necessary, and the classification evidence should be easily exhibited for peer review. Taken above concerns into consideration, we developed VCE to classify variants identified in our study. It could perform the classification automatically based on objective evidence and mark variants that need manual check. It is also open to the further extend of literature database, so that it will become more applicable and less manual intervention will be needed.

There is a breast cancer cumulative risk of people carrying a *BRCA1*, *BRCA2* or *PALB2* pathogenic variation (69, 70). Additional hereditary breast cancer-related susceptibility genes have also been studied by multiple gene panel sequencing previously(37). However, only a limited number of such studies have been conducted in China, due to the high cost of massive sequencing. Our data demonstrates a pathogenic mutation frequency in BRCA1 and BRCA2 of 11%, similar to other studies. Another large Chinese study was published from Shanghai, reporting a 9.1% frequency in women with at least one breast cancer risk factor(14). They identified the BRCA1 c.5470_5477del as the common mutation in their population, a mutation we also report as a recurrent mutation. Besides BRCA1 and BRCA2, PALB2 and ATM also showed high prevalence in Xie et al ‘s research(11) based on unselected breast cancer patient. HMMR and BRAD1 shows higher positive rate which remind those two genes may be important for high risk patients. In our cohort, the total positive rate of breast cancer related genes is 15.4%, a median rate between familial breast cancer and early onset breast cancer, and patients with age less than 40 is not associate with BRCA1/2 mutation carrier. These results car in line with the conclusion from Shao et al ‘s research(14) which indicates the cut off value may be set to 45.

The susceptibility genes of other cancer are mainly about gastric cancer and acute myeloid leukemia (AML). Those patients are enrolled in our research mainly because of their family history, and our clinical follow up also confirm that susceptibility genes carriers indeed may inherit the related familial diseases. Therefore, a comprehensive clinical genetic consulting including multiple department is needed. Otherwise the patient’s susceptibility genes of other cancer may be not recognized and well-treated. Another problem in variants interpretation is VUS. In our cohort, a total of 2632 VUS spots on 115 genes were identified in 787 out of 829 subjects, among which 684(82.5%) patients carried more than one VUS. According to other data from 1112 Shenzhen HBC high risk people, VUS rate of 21 breast cancer related gene is 55.04%. BRCA1 and BRCA2 in our data had (5.1%) and (12.1%) VUS rate respectively. Therefore, Chinese cohorts generally present a higher VUS rate than data from Myraid(71). It would be important to lower the frequencies of VUSs in Chinese cohorts. Recently, a large scale of variants validation on BRCA1 is conducted(67). This research provided new evidence for RING and BRCT domain in BRCA1, but there are still some short comes when applying it in clinical. For instance, in the experiments validating the VUS function, all the editing was conducted in the same cell line. However, in the real patient related situation, there will be a more complicate genetic background. It’s critical to confirm if those validation is consistent with clinical observation.

Our research is a multiple-gene test based hereditary breast cancer research with a big data set in China, which provides a comprehensive perspective of the breast cancer pathogenic variation in susceptibility gene as well as VUS spectrum in Chinese. With our data, we present the variant spectrum of 115 hereditary cancer-related susceptibility genes in 829 high-risk breast cancer patients.

In this research, we develop a semi-automatic variation classification tools and identified the function of five BRCA1 VUS in Chinese cohort. By analyzing our data, we provide a comprehensive overview of the pathogenic variant detection rate and the difference between Chinese and other populations. We have established screening pipeline and explorer the usage of multiple-gene hereditary cancer test in clinical practice of China. This work will aid the prevention and treatment of breast cancer in the Chinese people.

## Abbreviations

NCCN: National Comprehensive Cancer Network
HGMD: Human Gene Mutation Database
VUS: variants of uncertain significance
ACMG: American College of Medical Genetics

## Declarations

## Acknowledgement

Our research was supported by the following grants: National Natural Science Foundation of China(81172498/H1622, U1301252); Project of Heilongjiang province applied technology research and development(GA13C201); National Key Technology Support Program(2013BAI09B00); Specific research fund for public service sector, national health and family planning commission of the People’s Republic of China (201402003), National Key R&D Program of China (2017YFC1309103), the Science, Technology and Innovation Committee of Shenzhen Municipality (JSGG20140702161347218, CXZZ20140414170821163), Shenzhen Engineering Laboratory for Innovative Molecular Diagnostics [grant number DRC-SZ [2016] 884] funded by Development and Reform Commission of Shenzhen Municipality and by the intramural research program of the Division of Cancer Epidemiology and Genetics, National Cancer Institute, National Institutes of Health.

## Authors’ contributions

Conception and direction: Da Pang, Guibo Li, Shida Zhu, Yong Hou, Xianming Wang, Michael Dean, Xiaofei Ye, Huanming Yang, Jian Wang.

Acquisition of pedigree information and blood: Xianyu Zhang, Bingbing Song, Min Wang, Wenjing Jian, Jingjing Xie, Bingshu Xia, Shouping Xu.

Sample treatment, sequencing and data analysis: Xiaohong Wang, Boyang Cao, Kang Shao, Meng Liu, Liyun Xiao, Zhao Zhang, Enhao Fang, Haoxuan Jin, Xiaofeng Wei.

Manuscript writing: Xiaohong Wang, Kang Shao, Michael Dean, Cong Lin

## Supplementary

### Comparison of the two sequencing platforms

DNA was extracted from YH cells - a normal cell line, used to construct a library, captured by the same BGI Chip, and sequenced on the two different platforms, Blackbird and Hiseq 2500, respectively. To compare the divergence in variant calls on these sequencing platforms, 0.6G base data were generated with at least 400X and 200X depth by Blackbird and Hiseq 2500 platform respectively, and 99% coverage on target regions. Variants were then called by the corresponding pipelines. In total, 409 SNPs and 38 Indels were identified on both the Blackbird and Hiseq 2500 platforms; 8 SNPs and 7 Indels were specifically detected on the Blackbird platform. 36 SNPs and 51 indels were called on Hiseq 2500 platform specifically. Then all the SNPs, all 7 indels detected by Blackbird and 28 of 51 indels detected by Hiseq were selected for sanger sequencing.

Sanger sequencing was used to validate selected variants from these sequencing platforms. Pairs of primers for amplification and Sanger sequencing were designed to target regions flanking 29 out of 44 SNPs and 32 out of 35 Indels. The rest of the variants were unable to be confirmed due to their location in repeat regions. PCR amplification and Sanger sequencing were then performed. On the Blackbird platform, 7 SNPs and 2 Indels were validated, 1 Indel was not, and 4 Indels were unable to be confirmed due to the duplication of A or T. However on the Hiseq 2500 platform, 12 SNPs and 3 Indels were both validated, 2 SNPs and 2 Indels not, and 8 SNPs and 20 Indels unable to call.(Supplementary Figure S1 and Table S3).

## Reference

1. Narod SA, Iqbal J, Miller AB. Why have breast cancer mortality rates declined? Journal of Cancer Policy. 2015;5:8–17.

2. Chen W, Zheng R, Baade PD, Zhang S, Zeng H, Bray F, et al. Cancer statistics in China, 2015. CA: a cancer journal for clinicians. 2016;66:115–32.

3. Hall MJ, Reid JE, Burbidge LA, Pruss D, Deffenbaugh AM, Frye C, et al. BRCA1 and BRCA2 mutations in women of different ethnicities undergoing testing for hereditary breast-ovarian cancer. Cancer. 2009;115:2222–33.

4. King MC, Levy-Lahad E, Lahad A. Population-based screening for BRCA1 and BRCA2: 2014 Lasker Award. Jama. 2014;312:1091–2.

5. Kauff ND, Satagopan JM, Robson ME, Scheuer L, Hensley M, Hudis CA, et al. Risk-reducing salpingo-oophorectomy in women with a BRCA1 or BRCA2 mutation. N Engl J Med. 2002;346:1609–15.

6. Gabai-Kapara E, Lahad A, Kaufman B, Friedman E, Segev S, Renbaum P, et al. Population-based screening for breast and ovarian cancer risk due to BRCA1 and BRCA2. Proceedings of the National Academy of Sciences of the United States of America. 2014;111:14205–10.

7. Desmond A, Kurian AW, Gabree M, Mills MA, Anderson MJ, Kobayashi Y, et al. Clinical Actionability of Multigene Panel Testing for Hereditary Breast and Ovarian Cancer Risk Assessment. JAMA Oncol. 2015;1:943–51.

8. Dean M, Boland J, Yeager M, Im KM, Garland L, Rodriguez-Herrera M, et al. Addressing health disparities in Hispanic breast cancer: accurate and inexpensive sequencing of BRCA1 and BRCA2. Gigascience. 2015;4:50.

9. Dutil J, Golubeva VA, Pacheco-Torres AL, Diaz-Zabala HJ, Matta JL, Monteiro AN. The spectrum of BRCA1 and BRCA2 alleles in Latin America and the Caribbean: a clinical perspective. Breast Cancer Res Treat. 2015;154:441–53.

10. Tung N, Lin NU, Kidd J, Allen BA, Singh N, Wenstrup RJ, et al. Frequency of Germline Mutations in 25 Cancer Susceptibility Genes in a Sequential Series of Patients With Breast Cancer. Journal of clinical oncology: official journal of the American Society of Clinical Oncology. 2016;34:1460–8.

11. Sun J, Meng H, Yao L, Lv M, Bai J, Zhang J, et al. Germline Mutations in Cancer Susceptibility Genes in a Large Series of Unselected Breast Cancer Patients. Clinical cancer research: an official journal of the American Association for Cancer Research. 2017;23:6113–9.

12. Yang XR, Devi BCR, Sung H, Guida J, Mucaki EJ, Xiao Y, et al. Prevalence and spectrum of germline rare variants in BRCA1/2 and PALB2 among breast cancer cases in Sarawak, Malaysia. Breast Cancer Res Treat. 2017.

13. Wen WX, Allen J, Lai KN, Mariapun S, Hasan SN, Ng PS, et al. Inherited mutations in BRCA1 and BRCA2 in an unselected multiethnic cohort of Asian patients with breast cancer and healthy controls from Malaysia. J Med Genet. 2017.

14. Lang GT, Shi JX, Hu X, Zhang CH, Shan L, Song CG, et al. The spectrum of BRCA mutations and characteristics of BRCA-associated breast cancers in China: Screening of 2,991 patients and 1,043 controls by next-generation sequencing. International journal of cancer. 2017;141:129–42.

15. Seong MW, Cho S, Noh DY, Han W, Kim SW, Park CM, et al. Comprehensive mutational analysis of BRCA1/BRCA2 for Korean breast cancer patients: evidence of a founder mutation. Clin Genet. 2009;76:152–60.

16. Emi M, Matsushima M, Katagiri T, Yoshimoto M, Kasumi F, Yokota T, et al. Multiplex mutation screening of the BRCA1 gene in 1000 Japanese breast cancers. Jpn J Cancer Res. 1998;89:12–6.

17. Haitian Z, Yunfei L, Jian Z, Jian L, Qinghua L, Fuqiang W. Mutation screening of the BRCA1 gene in sporadic breast cancer in southern Chinese populations. Breast. 2008;17:563–7.

18. Li G, Guo X, Tang L, Chen M, Luo X, Peng L, et al. Analysis of BRCA1/2 mutation spectrum and prevalence in unselected Chinese breast cancer patients by next-generation sequencing. J Cancer Res Clin Oncol. 2017;143:2011–24.

19. Guan Y, Hu H, Peng Y, Gong Y, Yi Y, Shao L, et al. Detection of inherited mutations for hereditary cancer using target enrichment and next generation sequencing. Familial cancer. 2015;14:9.

20. Richards CS, Bale S, Bellissimo DB, Das S, Grody WW, Hegde MR, et al. ACMG recommendations for standards for interpretation and reporting of sequence variations: Revisions 2007. Genetics in medicine: official journal of the American College of Medical Genetics. 2008;10:294–300.

21. Richards S, Aziz N, Bale S, Bick D, Das S, Gastier-Foster J, et al. Standards and guidelines for the interpretation of sequence variants: a joint consensus recommendation of the American College of Medical Genetics and Genomics and the Association for Molecular Pathology. Genetics in medicine: official journal of the American College of Medical Genetics. 2015;17:405–24.

22. Starita LM, Young DL, Islam M, Kitzman JO, Gullingsrud J, Hause RJ, et al. Massively Parallel Functional Analysis of BRCA1 RING Domain Variants. Genetics. 2015;200:413–22.

23. Humphrey JS, Salim A, Erdos MR, Collins FS, Brody LC, Klausner RD. Human BRCA1 inhibits growth in yeast: potential use in diagnostic testing. Proceedings of the National Academy of Sciences of the United States of America. 1997;94:5820–5.

24. Li Q, Wang K. InterVar: Clinical Interpretation of Genetic Variants by the 2015 ACMG-AMP Guidelines. American journal of human genetics. 2017;100:267–80.

25. Scott AD, Huang KL, Weerasinghe A, Mashl RJ, Gao Q, Martins Rodrigues F, et al. CharGer: Clinical Characterization of Germline Variants. Bioinformatics. 2018.

26. Lin PH, Kuo WH, Huang AC, Lu YS, Lin CH, Kuo SH, et al. Multiple gene sequencing for risk assessment in patients with early-onset or familial breast cancer. Oncotarget. 2016;7:8310–20.

27. Yang X, Wu J, Lu J, Liu G, Di G, Chen C, et al. Identification of a comprehensive spectrum of genetic factors for hereditary breast cancer in a Chinese population by next-generation sequencing. PLoS One. 2015;10:e0125571.

28. Kwong A, Ng EK, Wong CL, Law FB, Au T, Wong HN, et al. Identification of BRCA1/2 founder mutations in Southern Chinese breast cancer patients using gene sequencing and high resolution DNA melting analysis. PLoS One. 2012;7:e43994.

29. Zhang J, Pei R, Pang Z, Ouyang T, Li J, Wang T, et al. Prevalence and characterization of BRCA1 and BRCA2 germline mutations in Chinese women with familial breast cancer. Breast Cancer Res Treat. 2012;132:421–8.

30. Vaidyanathan K, Lakhotia S, Ravishankar HM, Tabassum U, Mukherjee G, Somasundaram K. BRCA1 and BRCA2 germline mutation analysis among Indian women from south India: identification of four novel mutations and high-frequency occurrence of 185delAG mutation. Journal of biosciences. 2009;34:415–22.

31. Sugano K, Nakamura S, Ando J, Takayama S, Kamata H, Sekiguchi I, et al. Cross-sectional analysis of germline BRCA1 and BRCA2 mutations in Japanese patients suspected to have hereditary breast/ovarian cancer. Cancer Science. 2008;99:1967–76.

32. Ikeda N, Miyoshi Y, Yoneda K, Shiba E, Sekihara Y, Kinoshita M, et al. Frequency of BRCA1 and BRCA2 germline mutations in Japanese breast cancer families. International journal of cancer. 2001;91:83–8.

33. Ang P, Lim IH, Lee TC, Luo JT, Ong DC, Tan PH, et al. BRCA1 and BRCA2 mutations in an Asian clinic-based population detected using a comprehensive strategy. Cancer Epidemiol Biomarkers Prev. 2007;16:2276–84.

34. Rashid MU, Zaidi A, Torres D, Sultan F, Benner A, Naqvi B, et al. Prevalence of BRCA1 and BRCA2 mutations in Pakistani breast and ovarian cancer patients. International journal of cancer. 2006;119:2832–9.

35. Llort G, Munoz CY, Tuser MP, Guillermo IB, Lluch JR, Bale AE, et al. Low frequency of recurrent BRCA1 and BRCA2 mutations in Spain. Hum Mutat. 2002;19:307.

36. Torres D, Rashid MU, Gil F, Umana A, Ramelli G, Robledo JF, et al. High proportion of BRCA1/2 founder mutations in Hispanic breast/ovarian cancer families from Colombia. Breast Cancer Res Treat. 2007;103:225–32.

37. Kurian AW, Hare EE, Mills MA, Kingham KE, McPherson L, Whittemore AS, et al. Clinical evaluation of a multiple-gene sequencing panel for hereditary cancer risk assessment. Journal of clinical oncology: official journal of the American Society of Clinical Oncology. 2014;32:2001–9.

38. Tung N, Battelli C, Allen B, Kaldate R, Bhatnagar S, Bowles K, et al. Frequency of mutations in individuals with breast cancer referred for BRCA1 and BRCA2 testing using next-generation sequencing with a 25-gene panel. Cancer. 2015;121:25–33.

39. Zhi X, Szabo C, Chopin S, Suter N, Wang QS, Ostrander EA, et al. BRCA1 and BRCA2 sequence variants in Chinese breast cancer families. Hum Mutat. 2002;20:474.

40. Li WF, Hu Z, Rao NY, Song CG, Zhang B, Cao MZ, et al. The prevalence of BRCA1 and BRCA2 germline mutations in high-risk breast cancer patients of Chinese Han nationality: two recurrent mutations were identified. Breast Cancer Res Treat. 2008;110:99–109.

41. Kim H, Cho DY, Choi DH, Choi SY, Shin I, Park W, et al. Characteristics and spectrum of BRCA1 and BRCA2 mutations in 3,922 Korean patients with breast and ovarian cancer. Breast Cancer Res Treat. 2012;134:1315–26.

42. Cao WM, Gao Y, Yang HJ, Xie SN, Ding XW, Pan ZW, et al. Novel germline mutations and unclassified variants of BRCA1 and BRCA2 genes in Chinese women with familial breast/ovarian cancer. BMC cancer. 2016;16:64.

43. Choi DH, Lee MH, Bale AE, Carter D, Haffty BG. Incidence of BRCA1 and BRCA2 mutations in young Korean breast cancer patients. Journal of clinical oncology: official journal of the American Society of Clinical Oncology. 2004;22:1638–45.

44. Plaschke J, Commer T, Jacobi C, Schackert HK, Chang-Claude J. BRCA2 germline mutations among early onset breast cancer patients unselected for family history of the disease. J Med Genet. 2000;37:E17.

45. Bergthorsson JT, Ejlertsen B, Olsen JH, Borg A, Nielsen KV, Barkardottir RB, et al. BRCA1 and BRCA2 mutation status and cancer family history of Danish women affected with multifocal or bilateral breast cancer at a young age. J Med Genet. 2001;38:361–8.

46. Nielsen HR, Nilbert M, Petersen J, Ladelund S, Thomassen M, Pedersen IS, et al. BRCA1/BRCA2 founder mutations and cancer risks: impact in the western Danish population. Fam Cancer. 2016;15:507–12.

47. Wu Y, Berends MJ, Sijmons RH, Mensink RG, Verlind E, Kooi KA, et al. A role for MLH3 in hereditary nonpolyposis colorectal cancer. Nature genetics. 2001;29:137–8.

48. Tischkowitz MD, Morgan NV, Grimwade D, Eddy C, Ball S, Vorechovsky I, et al. Deletion and reduced expression of the Fanconi anemia FANCA gene in sporadic acute myeloid leukemia. Leukemia. 2004;18:420–5.

49. Kee Y, D’Andrea AD. Molecular pathogenesis and clinical management of Fanconi anemia. The Journal of clinical investigation. 2012;122:3799–806.

50. Stieglitz E, Loh ML. Genetic predispositions to childhood leukemia. Therapeutic advances in hematology. 2013;4:270–90.

51. Boocock GR, Morrison JA, Popovic M, Richards N, Ellis L, Durie PR, et al. Mutations in SBDS are associated with Shwachman-Diamond syndrome. Nature genetics. 2003;33:97–101.

52. Hall GW, Dale P, Dodge JA. Shwachman-Diamond syndrome: UK perspective. Archives of disease in childhood. 2006;91:521–4.

53. Ihara K, Ishii E, Eguchi M, Takada H, Suminoe A, Good RA, et al. Identification of mutations in the c-mpl gene in congenital amegakaryocytic thrombocytopenia. Proceedings of the National Academy of Sciences of the United States of America. 1999;96:3132–6.

54. Germeshausen M, Ballmaier M, Welte K. MPL mutations in 23 patients suffering from congenital amegakaryocytic thrombocytopenia: the type of mutation predicts the course of the disease. Hum Mutat. 2006;27:296.

55. Roa BB, Savino CV, Richards CS. Ashkenazi Jewish population frequency of the Bloom syndrome gene 2281 delta 6ins7 mutation. Genetic testing. 1999;3:219–21.

56. German J. Bloom’s syndrome. XX. The first 100 cancers. Cancer genetics and cytogenetics. 1997;93:100–6.

57. Gimm O, Greco A, Hoang-Vu C, Dralle H, Pierotti MA, Eng C. Mutation analysis reveals novel sequence variants in NTRK1 in sporadic human medullary thyroid carcinoma. The Journal of clinical endocrinology and metabolism. 1999;84:2784–7.

58. Clementi R, Locatelli F, Dupre L, Garaventa A, Emmi L, Bregni M, et al. A proportion of patients with lymphoma may harbor mutations of the perforin gene. Blood. 2005;105:4424–8.

59. El Abed R, Bourdon V, Voskoboinik I, Omri H, Youssef YB, Laatiri MA, et al. Molecular study of the perforin gene in familial hematological malignancies. Hereditary cancer in clinical practice. 2011;9:9.

60. Chaudhry MS, Gilmour KC, House IG, Layton M, Panoskaltsis N, Sohal M, et al. Missense mutations in the perforin (PRF1) gene as a cause of hereditary cancer predisposition. Oncoimmunology. 2016;5:e1179415.

61. Friedman JM. Epidemiology of neurofibromatosis type 1. American journal of medical genetics. 1999;89:1–6.

62. Williams VC, Lucas J, Babcock MA, Gutmann DH, Korf B, Maria BL. Neurofibromatosis type 1 revisited. Pediatrics. 2009;123:124–33.

63. Kitao S, Shimamoto A, Goto M, Miller RW, Smithson WA, Lindor NM, et al. Mutations in RECQL4 cause a subset of cases of Rothmund-Thomson syndrome. Nature genetics. 1999;22:82–4.

64. Wang LL, Levy ML, Lewis RA, Chintagumpala MM, Lev D, Rogers M, et al. Clinical manifestations in a cohort of 41 Rothmund-Thomson syndrome patients. American journal of medical genetics. 2001;102:11–7.

65. Aoki Y, Niihori T, Kawame H, Kurosawa K, Ohashi H, Tanaka Y, et al. Germline mutations in HRAS proto-oncogene cause Costello syndrome. Nature genetics. 2005;37:1038–40.

66. Sun X, Frierson HF, Chen C, Li C, Ran Q, Otto KB, et al. Frequent somatic mutations of the transcription factor ATBF1 in human prostate cancer. Nature genetics. 2005;37:407–12.

67. Findlay GM, Daza RM, Martin B, Zhang MD, Leith AP, Gasperini M, et al. Accurate classification of BRCA1 variants with saturation genome editing. Nature. 2018;562:217–22.

68. Amendola LM, Jarvik GP, Leo MC, McLaughlin HM, Akkari Y, Amaral MD, et al. Performance of ACMG-AMP Variant-Interpretation Guidelines among Nine Laboratories in the Clinical Sequencing Exploratory Research Consortium. American journal of human genetics. 2016;98:1067–76.

69. Ford D, Easton DF, Stratton M, Narod S, Goldgar D, Devilee P, et al. Genetic heterogeneity and penetrance analysis of the BRCA1 and BRCA2 genes in breast cancer families. The Breast Cancer Linkage Consortium. American journal of human genetics. 1998;62:676–89.

70. Antoniou AC, Casadei S, Heikkinen T, Barrowdale D, Pylkas K, Roberts J, et al. Breast-cancer risk in families with mutations in PALB2. N Engl J Med. 2014;371:497–506.

71. Eggington JM, Bowles KR, Moyes K, Manley S, Esterling L, Sizemore S, et al. A comprehensive laboratory-based program for classification of variants of uncertain significance in hereditary cancer genes. Clin Genet. 2014;86:229–37.

