## supplementary material figure1 and table3 for "Gene Panel Sequencing in a Chinese High-risk Breast Cancer Cohort"

A	B

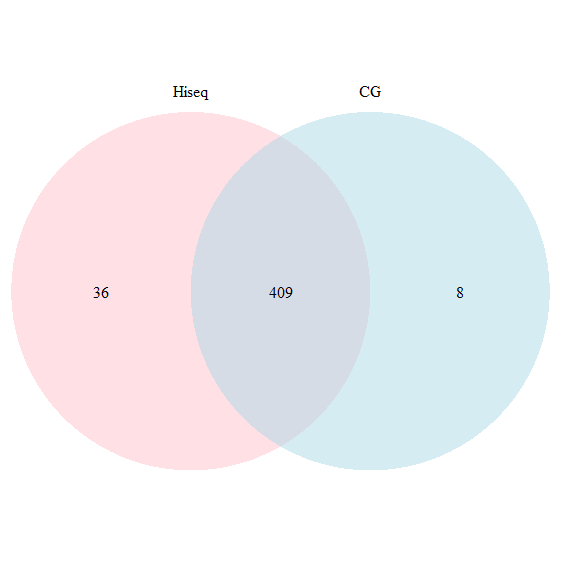

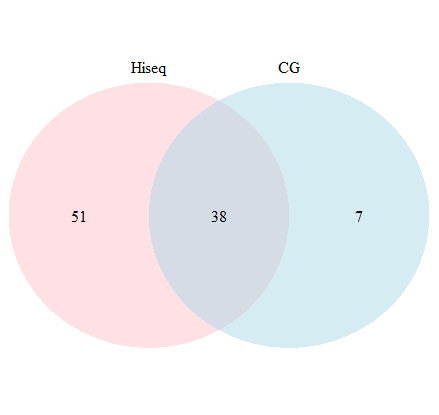

Fig S1 A. SNPs called by different sequencing platforms, B. Indels called by different sequencing platforms

Table S3. Verification results by Sanger sequencing

| SNP/Indel | True/false | Blackbird | Hiseq 2500 |
| --- | --- | --- | --- |
| SNP | true | 7/7 | 12/22 |
|  | false | 0/7 | 2/22 |
| Indel | true | 2/7 | 3/25 |
|  | false | 1/7 | 2/25 |
